# A Multimodal and Integrated Approach to Interrogate Human Kidney Biopsies with Rigor and Reproducibility: The Kidney Precision Medicine Project

**DOI:** 10.1101/828665

**Authors:** Tarek M. El-Achkar, Michael T. Eadon, Rajasree Menon, Blue B. Lake, Tara K. Sigdel, Theodore Alexandrov, Samir Parikh, Guanshi Zhang, Dejan Dobi, Kenneth W. Dunn, Edgar A. Otto, Christopher R. Anderton, Jonas M. Carson, Jinghui Luo, Chris Park, Habib Hamidi, Jian Zhou, Paul Hoover, Andrew Schroeder, Marianinha Joanes, Evren U. Azeloglu, Rachel Sealfon, Seth Winfree, Becky Steck, Yongqun He, Vivette D’Agati, Ravi Iyengar, Olga G Troyanskaya, Laura Barisoni, Joseph Gaut, Kun Zhang, Zoltan Laszik, Brad Rovin, Pierre C. Dagher, Kumar Sharma, Minnie Sarwal, Jeffrey B. Hodgin, Charles E. Alpers, Matthias Kretzler, Sanjay Jain, For the Kidney Precision Medicine Project

## Abstract

Comprehensive and spatially mapped molecular atlases of organs at a cellular level are a critical resource to gain insights into pathogenic mechanisms and personalized therapies for diseases. The Kidney Precision Medicine Project (KPMP) is an endeavor to generate 3-dimensional (3D) molecular atlases of healthy and diseased kidney biopsies using multiple state-of-the-art OMICS and imaging technologies across several institutions. Obtaining rigorous and reproducible results from disparate methods and at different sites to interrogate biomolecules at a single cell level or in 3D space is a significant challenge that can be a futile exercise if not well controlled. We describe a “follow the tissue” pipeline for generating a reliable and authentic single cell/region 3D molecular atlas of human adult kidney. Our approach emphasizes quality assurance, quality control, validation and harmonization across different OMICS and imaging technologies from sample procurement, processing, storage, shipping to data generation, analysis and sharing. We established benchmarks for quality control, rigor, reproducibility and feasibility across multiple technologies through a pilot experiment using common source tissue that was processed and analyzed at different institutions and different technologies. A peer review system was established to critically review quality control measures and the reproducibility of data generated by each technology before being approved to interrogate clinical biopsy specimens. The process established economizes the use of valuable biopsy tissue for multi-OMICS and imaging analysis with stringent quality control to ensure rigor and reproducibility of results and serves as a model for precision medicine projects across laboratories, institutions and consortia.

## Introduction

Recent advances in biotechnology allow capturing the state of a tissue in health and disease at an unprecedented structural and molecular resolution.(9) Application of these technologies at the level of the genome, transcriptome, proteome and metabolome have enabled identification of regulatory cascades and their mapping into tissue compartments at a single cell resolution.(12, 17, 18, 22, 23, 25, 27) There is an urgent need to apply these technologies to clinical samples from patients with the two most devastating categories of kidney diseases, chronic kidney disease (CKD) and acute kidney injury (AKI). With a prevalence as high as 14% (37 million people) for CKD and high mortality in AKI patients, deciphering the underlying molecular and architectural complexity can result in better treatment in these conditions.(20) Investigators have begun to apply single cell OMICS and high resolution imaging technologies to healthy or diseased kidney biopsy tissue to provide important information on the cell type composition and spatial relationships in diseases like lupus nephritis, diabetes and in healthy tissue.(21, 24) Integrating multimodal information derived from existing datasets and emerging technologies is a major challenge, because protocols, biological terms and experimental standards are not uniform. In addition, applying multiple cutting-edge technologies frequently involve the collaboration of several labs with specialized practices and protocols.

Recognizing these limitations, one of the goals of the Kidney Precision Medicine Project (KPMP) is to establish rigorous pre-analytical and analytical protocols with highly standardized and controlled workflows to interrogate biopsies of AKI and CKD patients using cutting edge OMICS and imaging technologies. Here we describe a quality-controlled tissue interrogation pipeline component of the KPMP for multimodal analysis of kidney biopsies. This pipeline realizes the power of combined analyses of well-vetted and curated data from different technologies to ensure rigor, reproducibility and complementarity to generate molecular atlases of healthy and disease kidneys that can impact patient care, while extracting maximum data from a limited amount of tissue. The framework established serves as a paradigm for similar atlas efforts and precision medicine projects of other organs systems and diseases and a guide to investigators to generate high quality and reproducible data from limited tissue.(1, 9, 10, 17)

## Results and Discussion

### Guiding principles for multimodal quality-controlled tissue interrogation

The KPMP encompasses a diverse set of technologies to generate a robust molecular, spatial and structural atlas (**Fig.1, 2, Supplemental File 1**). A major strength of multimodal interrogation (**Table 1** lists specialized terms used) of various biomolecules including RNA, proteins and metabolites using different technologies is to ensure a comprehensive coverage of these biomolecules in case a single technology does not capture (“blind spots”) expression of a particular gene, protein or metabolite. Redundancy among the different technologies further provides orthogonal validation that lends confidence to the discovered genes/proteins/metabolites/cell types and cell states. To enhance data quality, reproducibility and identify weaknesses and strengths of each technology (**Fig. 2** and see later) in this multimodal approach our guiding principle was to harmonize (**Table 1)** tissue collection, processing, preservation and analytical steps.

**Fig. 1.**
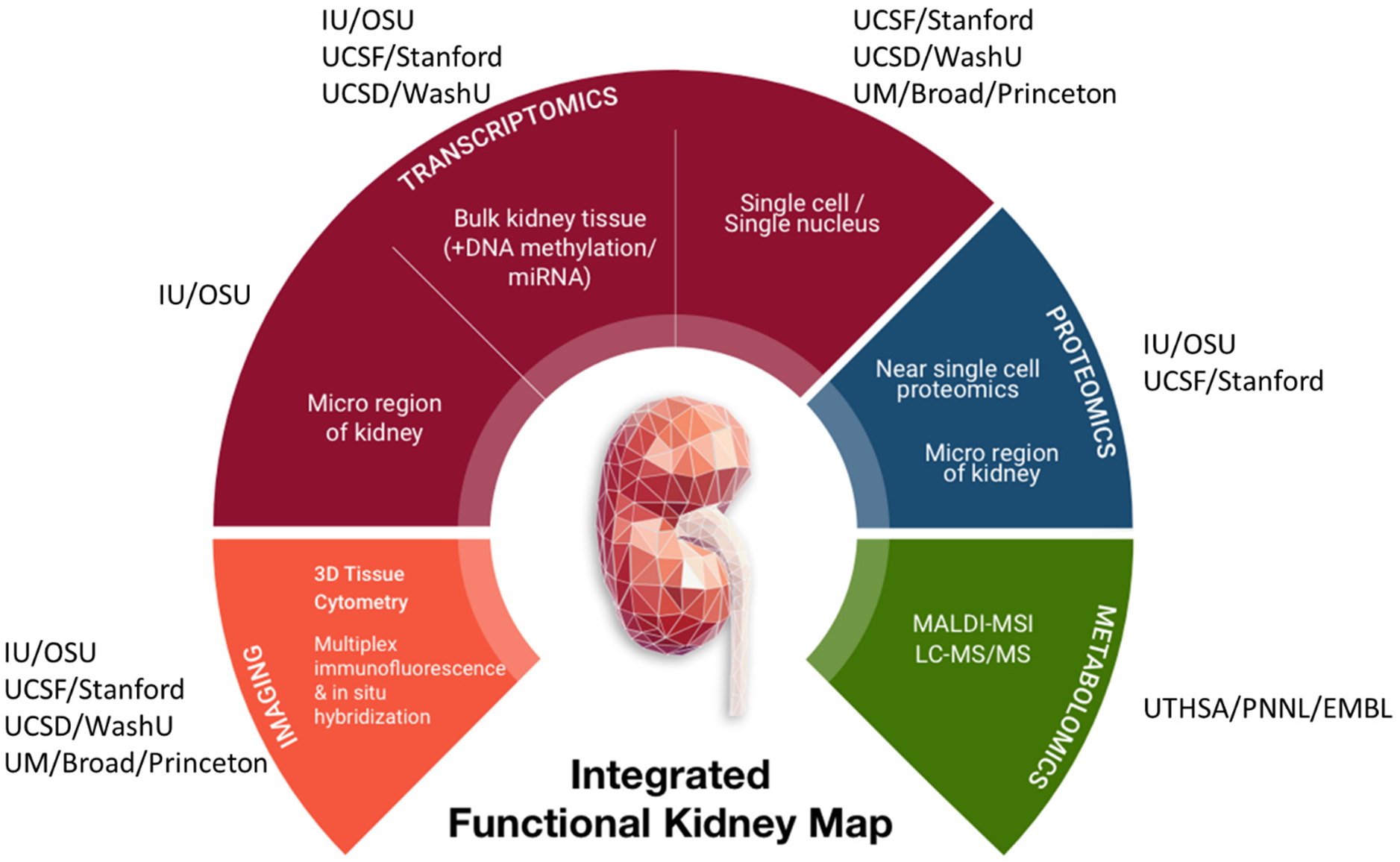
Overview of Tissue Interrogation Sites and technologies in KPMP. IU-Indiana University; OSU – Ohio State University; UCSD – University of California, San Diego; WU – Washington University in St. Louis; UM – University of Michigan; UCSF – University of California, San Francisco; UTHSA – University of Texas Health Sciences, San Antonio; PNNL – Pacific Northwest National Laboratory; EMBL – European Molecular Biology Laboratory.

**Fig. 2.**
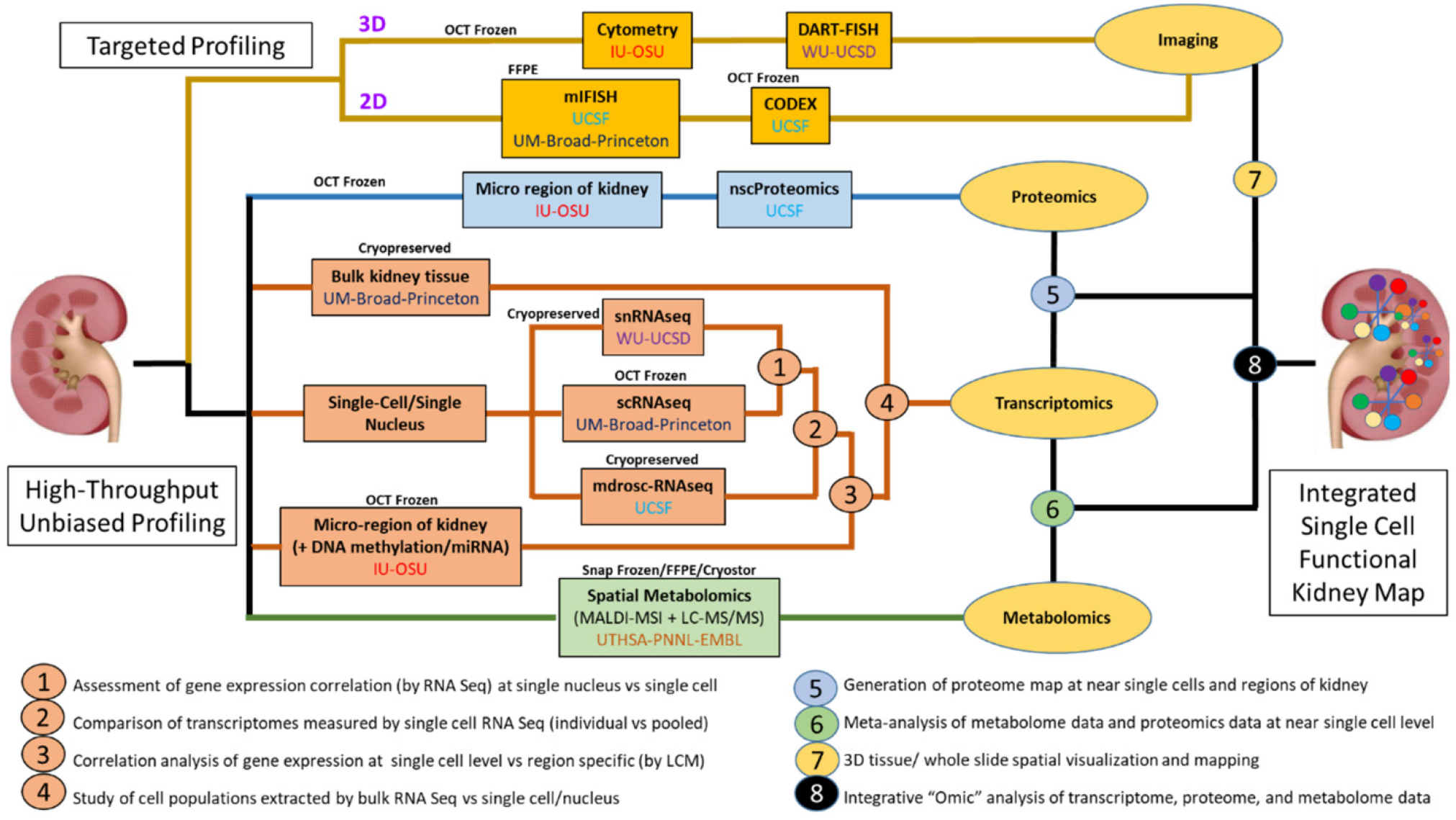
The blueprint for interconnectivity for multimodal molecular assessment of kidney tissue at the initiation of the KPMP.

**Table 1.** Description of selected quality control terms as they relate to the KPMP.

| **Term** | **Description** |
| --- | --- |
| Complementarity | Different types of information gained from different technologies to describe the tissue composition and the relationships among its constituents |
| Follow the tissue | The different steps that the tissue undergoes during its interrogation from collection to preservation, processing, storage, shipping, assay, analysis and data dissemination |
| Harmonization | Efforts to combine data from different sources including file formats, nomenclature into a cohesive and comparable format for analysis and interpretation |
| Metadata | Information or data associated with each aspect of the process of tissue collection, its interrogation and analysis |
| Metadata modeling | Development of rules and framework to define how to build relationships between concepts of a defined domain |
| Multimodal | Different ways (technologies) to interrogate the tissue in KPMP |
| Ontologize | The act of converting into ontological terms or entities |
| Ontology | A human- and computer-interpretable set of terms and relations that represent entities in a specific domain and how they relate to each other. |
| Orthogonal validation | Using different technologies or assays to verify a given observation. |
| Pipeline | The process through which each aspect of tissue interrogation is systematically done. Example, tissue preservation pipeline, tissue processing pipeline) |
| Standardization | The process by which critical elements of quality control parameters in “follow the tissue” are established and implemented to minimize sources of errors and technical variations and ensure consistent results are produced for a particular technology |
| Technology drift | Changes in a technology or its components over time that can impact the reproducibility of results |
| Tissue Interrogation | The act of using different techniques and tools to identify, analyze and determine the 2D/3D relationships between cells, their extracellular environment and their molecular components including gene, protein and metabolite expression |

However, applying multi-scalar technologies on a limited amount of tissue in a collaborative manner among the various KPMP tissue interrogation sites (TISs) posits unprecedented challenges. Errors are compounded as the data are dependent on multiple processes and steps that begin from specimen procurement to data generation and analysis. There are several sources of random and technical variations that confound the outcomes and impact biological reproducibility and interpretation of data. An additional challenge is maximizing the application of these sensitive, big data technologies to clinical biopsies with limited tissue available for research. We developed a process to standardize and harmonize (**Table 1)**, where possible, the entire pipeline from tissue procurement to analysis (termed follow the tissue) to overcome these challenges. Key factors considered were economizing tissue usage, maximizing preservation and use for multiple technologies, and documenting quality of intermediate steps with clearly defined quality assurance and control criteria. We implemented rigorous procedures for transparency, technical and biological reproducibility across multiple sources of tissue procurement, analysis and interpretation and standards for quality control (QC).

#### Overview of the strategy: “Follow the tissue”

Anticipating that the tissue would come from different recruitment sites, our approach was to develop a tissue-processing pipeline that can be easily implemented at the bedside and applicable to multiple state-of-the-art interrogation technologies. We required that each TIS demonstrates feasibility, reproducibility, applies rigorous analytics to their technology, develops processing methods that economizes tissue use and fosters integrated analysis and quality control measures in collaboration with the other TISs. The challenges in the “follow the tissue” process included variable standards in collecting metadata, procurement, processing, sample preparation, analytical parameters, analysis and data sharing and deposition in a repository for public access. To tackle these challenges, we formed working groups for tissue processing that included expertise in OMICS, imaging, and pathology. These groups addressed quality control measures across four main categories: 1) Participants, 2) Tissue, 3) Assays and analysis, 4) Data hub (**Fig. 3**). The objective was to identify critical parameters and data to be collected in each of the 4 categories that are important for standardization and QC. The outcomes of the meetings and decision process were documented at the KPMP management website in “Basecamp” (www.basecamp.com) to enable easy access to archived documents and meeting minutes for future reference. A critical aspect was formulating and capturing well-vetted relevant metadata in each of these categories to enable the interpretation of molecular discoveries among the underlying biological variations related to healthy and disease phenotypes (**Table 2)**. We established benchmarks for the entire process, and a clear approach for data visualization and sharing for various types of users. Once the overall vision or “blueprint” was established, the pipeline was pressure-tested with a pilot project using adult human kidney tissue (see later). We describe below the individual steps of this workflow.

**Fig. 3.**
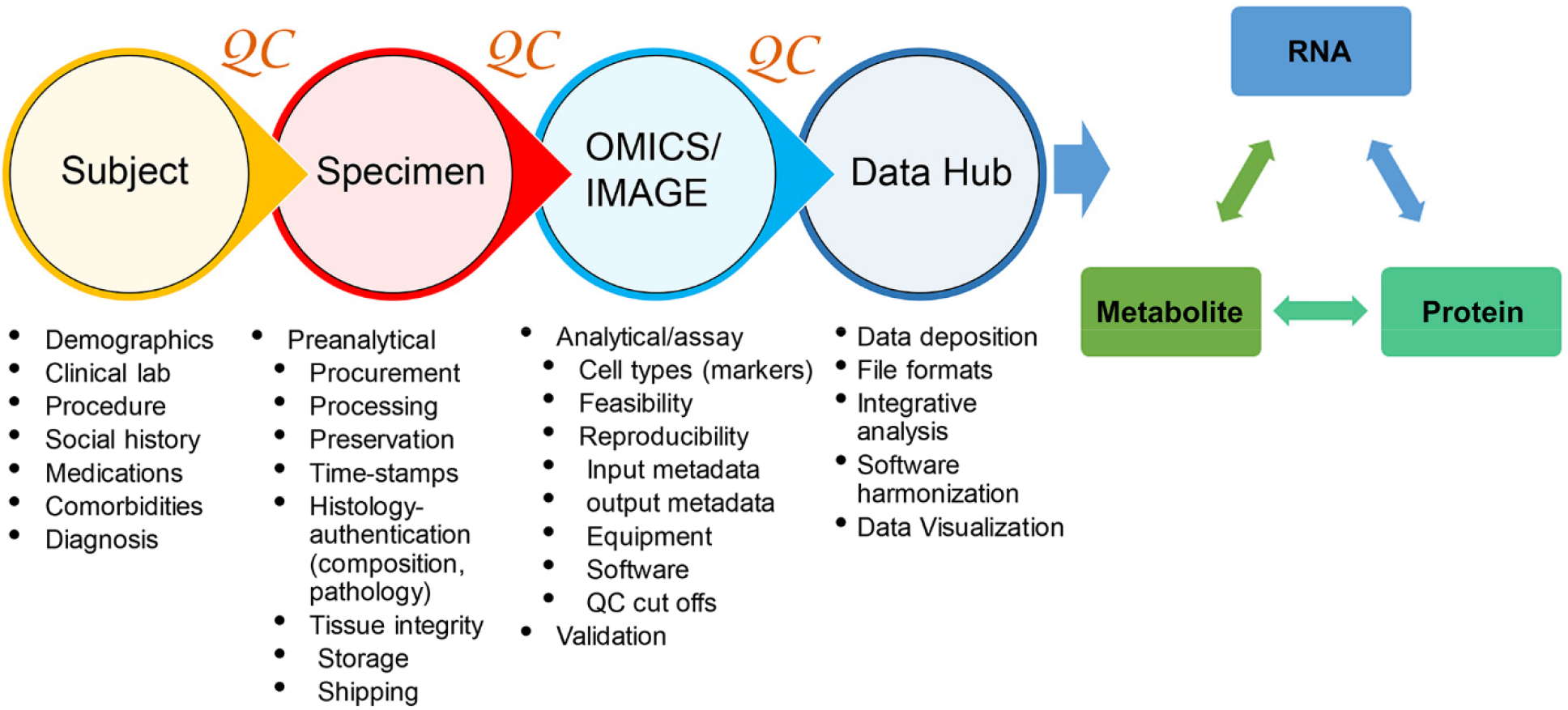
Overview of “Follow the Tissue” pipeline with essential QA/QC parameters.

**Table 2.** Preanalytical parameters for rigor, reproducibility, quality assessment and control.

| **Subject** | **Specimen Procurement** | **Specimen Morphology** | **Specimen Storage/Shipping** |
| --- | --- | --- | --- |
| Age | Procedure data | Gross assessment | Storage time |
| Race | Procedure Type (U/S, IR, Surgical) | Integrity (intact, fragmented) | Storage temperature |
| Sex | Location/Site | % Cortex | Storage medium |
| Height | Procurement time | % Medulla | Storage container |
| Weight | Specimen collection media | Microscopic assessment | Shipping time |
| BMI | Tissue transport time to lab | # Glomeruli | Shipping container |
| Clinical labs (blood, urine) | Transport temperature | % Global glomerulosclerosis (age corrected) | Shipping image |
| eGFR | Preservation media/condition | Necrosis | Receiving time |
| Clinical diagnosis | Supplies/Reagent details | Hemorrhage | Receiving image |
| Pathology diagnosis | Processing time | Autolysis | Storage time receiving |
| Comorbidities | Size | Tubular atrophy | Storage temp receiving |
| Medications | Image | Interstitial fibrosis |  |
| Social history |  | Interstitial inflammation |  |
| Family History |  | Image (gross, HE, PAS) |  |
|  |  | Freezing artifacts |  |

#### Subject Metadata

Detailed relevant parameters of participant associated metadata were identified through a team effort of all the recruitment sites and TISs and will be described elsewhere (**Table 2**). This was important to interpret variations in data due to contributions of patient attributes. For example, sex differences, age, race or medications such as diuretics or antihypertensives can contribute to changes in molecular or cellular distribution of transporters in the kidney. The metadata fields are being modeled and represented using the Kidney Tissue Atlas Ontology (KTAO),(7) and Ontology of Precision Medicine and Investigation (OPMI),(6) two open source biomedical ontologies that are being developed by collaborative efforts between the KPMP and ontology communities.

#### Tissue Preanalytical Considerations

We identified and harmonized several preanalytical parameters for QC related to specimen handling, processing, preservation, orientation, quantity used, shipping and storage to minimize their impact on technical variations and decreased data quality (**Table 2**). An infrastructure to track specimen movement from origin to specimen processing sites was developed (SpecTrack). A key feature was the ability to record deidentified specimens and documentation of the time, temperature and state in which the specimens were procured, shipped and received from the recruitment sites. All the sites were required to show successful use of this system prior to being qualified to receive specimens. An important consideration in designing the pipeline was to assess the quality and composition of the tissue being analyzed to best interpret the molecular outcomes. While some technologies depended on complete dissociation of tissue (scRNA-seq) others had the opportunity to register tissue composition before dissociation or analysis. As such, guidelines were developed by the KPMP pathologists and TIS investigators for high level quality assessment of composition and integrity of tissue sections and included relative proportions of cortex, medulla, glomerulosclerosis and features compromising integrity (**Table 2**).

#### Tissue Analytical and Assay Considerations

We focused on three main tenets: 1) Assay metadata, 2) Assay quality assurance (QA) parameters, and 3) Assay QC parameters. We emphasized from the beginning to define and record key metadata associated with assay performance to ensure transparency and reproducibility. QA is linked to understanding and applying the best practices recommended in the field for that particular technology, using a specific instrument set(s), protocols or platform. The reliance on data from the manufacturer or post-marketing analysis when available is essential. We ensured that all platforms used in data production are optimized to produce the best possible results or operating under *bone fide* core facilities. For the QC component, we expected to meet and exceed a set of criteria guaranteeing that the assay works properly. We harmonized metadata collections, assays, instrumentations and *post hoc* analyses for similar biomolecules and used standard terms where possible. QC parameters and minimum attributes that were relevant for the performance of the assay were identified and common terms were used for similar types of technologies (RNA or protein or imaging). Each TIS was expected to come up with concrete criteria for QC of each technology that could be tracked throughout and demonstrate pass-fail rates and reproducibility in pilot experiments (see later). Furthermore, within each technology, implementation of measures that allow detection and control of batch effects and assay drift were also incorporated. These criteria also set a benchmark to give reproducible data for building the kidney atlas (**Supplemental Tables S1-3**).

#### Identifying and annotating cell types

We followed the concept of building an iterative marker list derived from published data and data generated from the KPMP. This served several purposes: 1) To qualify the identity and composition of the tissue being interrogated, 2) To validate and optimize tissue processing pipelines, and 3) To identify regions or cell types for integrative quality check and analysis to build the kidney atlas. Our initial list (made in 2018) of a subset of cell type markers mainly relied on rodent studies and bulk RNAseq data with corresponding evidence from the human protein atlas (**Supplemental File 2).**(2–5, 13, 14, 19, 26, 28, 30) Later iterations of the potential cell types/states were heavily dependent on the data generated from the KPMP pilot project (see below). Similarly, for imaging studies a number of parameters were established to best standardize the formats of image acquisition, analysis and deposition (**Supplemental Table S4**).

#### Data quality check, visualization and sharing

After passing the local TIS QC the data were required to be deposited in the “data hub” that is managed by non-TIS members. The roles of the “data hub” team are: 1) the examination of the associated metadata for completeness, 2) independent analysis of the data for passing QC thresholds, 3) enhancing data availability to other sites of the KPMP for integrated analysis and quality check and 4) planning for public sharing. An essential component of data output is making it accessible to the public. In this regard, the KPMP data hub is tasked with a team dedicated for building tools for summary analysis and visualization of the integrated results generated by the various technologies.

### A consortium-wide pilot experiment to test the “follow the tissue” pipeline

#### Rationale

The objective of a consortium-wide pilot experiment was to use a same source kidney specimen to: 1) standardize tissue processing/handling, storage and shipping steps; 2) establish feasibility and validate the QA-QC parameters for all the technologies in the interrogation pipeline; 3) compare, when applicable across sites, the performance of molecular interrogation and identify sources of variabilities and concordance; 4) lay out a blueprint for harmonization and complementarity across technologies (**Table 1**); 5) identify gaps and weaknesses in the interrogation pipeline. An important outcome was to define a protocol that is harmonized across technologies, and that could ultimately be used for interrogating biopsies from patients in an economical and efficient manner.

#### Design

Tumor-free kidney cortical tissues from nephrectomy specimens were procured from the University of Michigan tissue collection center, preserved in different types of media according to the needs of the various TISs and distributed to each TIS for testing feasibility, validation and identifying the QC metrics for their respective technologies. Specifically, contiguous serial sections (approximately 1cm x 2mm x 2mm) in the shape of rectangular cuboids were cut for processing and preservation and shipped to each TIS designated by a code (**Fig. 4** and **Supplemental Fig. S1** for the preservation methods used). In total, 6 different nephrectomy specimens were processed as described above and used by all the TISs. Hence, not only each site had access to the same tissue source, but there were also 6 biological replicates distributed for the purpose of testing reproducibility, as discussed below.

**Fig. 4.**
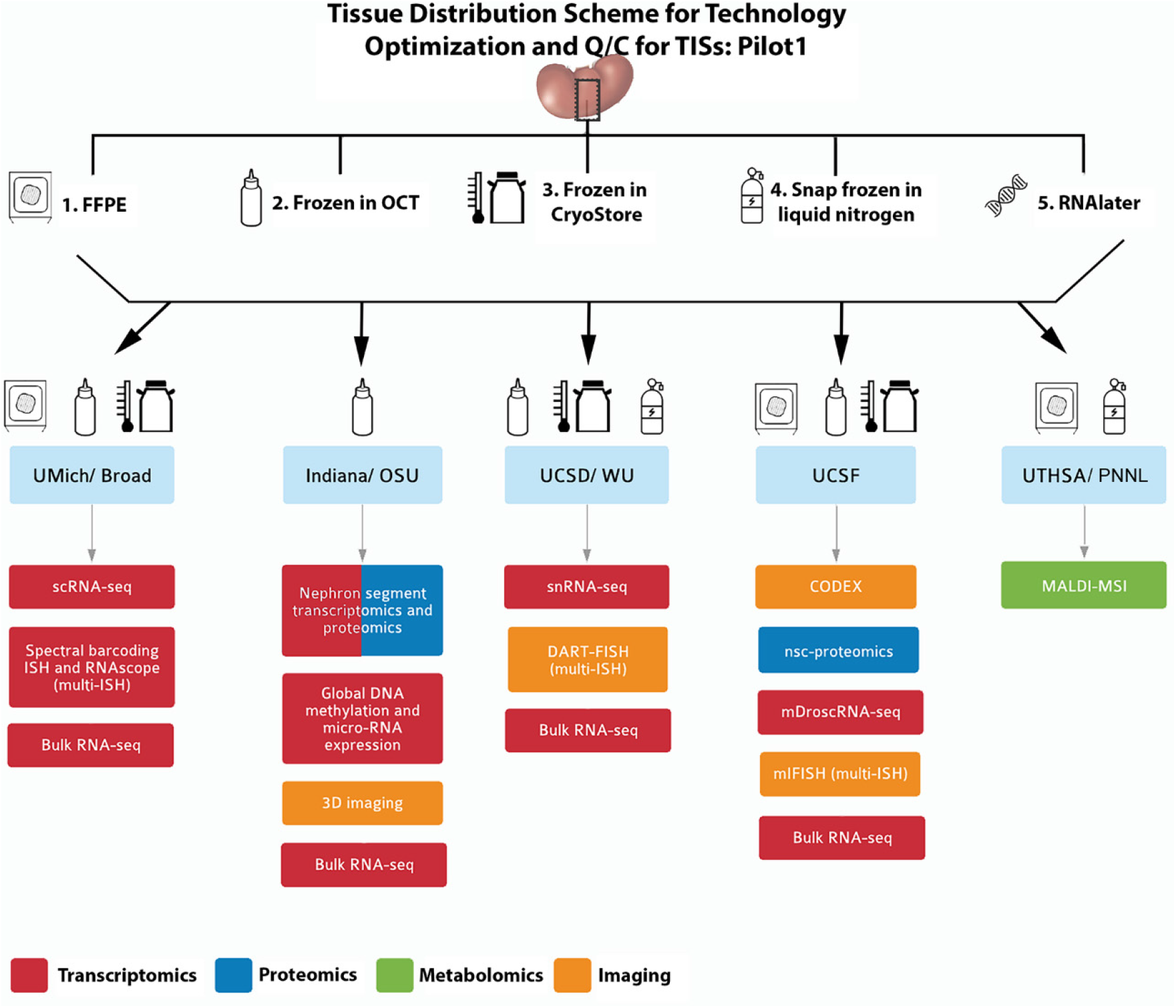
Pilot 1 tissue processing and distribution. A single source kidney tissue from a tumor nephrectomy was sectioned into equal blocks (schema in supplemental figure 1) and processed into 5 tissue processing methods according to the downstream applications listed. Blocks of the tissue was distributed to the sites according to the technology as indicated by the drawings. Formalin fixed paraffin embedded (FFPE); in-situ hybridization (ISH); single cell (sc) and single nuclear (sn) RNA sequencing (RNA-seq). Near single cell (nsc)-proteomics; matrix-assisted laser desorption/ionization mass spectrometry imaging (MALDI-MSI).

### Quality control outcomes and observations based on the pilot experiment

#### 1) Tissue procurement/preservation - Preanalytical parameters

The following outcomes were directly derived from this pilot experience:

A. *Better definition of the metadata associated with tissue procurement, preservation, integrity and composition.* We identified commonalities between tissue procurement, processing, assessment and storage that enabled the use of similar conditions for multiple technologies. For example, snRNAseq, 3D-tissue cytometry, LMD transcriptomics, LMD proteomics, mDroscRNAseq, spatial metabolomics, miFISH and DART-FISH could all use fresh frozen OCT blocks (**Supplemental Tables S1-3** and detailed in TIS manual of operations at www.kpmp.org/resources).
B. *Ontology-based metadata modeling and representation* (**Table 1**). The identified metadata elements confirmed the need to implement this ontology-based approach to represent relationships between different metadata types more meaningful and machine-interpretable, supporting advanced data analytics and knowledge discovery.(8, 15, 29)
C. *Real time testing of specimen tracking using the SpecTrack software.* This live tracking of the tissue revealed weaknesses in the pipeline and allowed improvements including better documentation of tissue and temperature states of shipments and appropriate packaging materials (**Supplemental Fig. 2,** see pathology protocols https://kpmp.org/researcher-resources/).
D. *Effect of shipping and best practices establishment.* To determine the effect of shipping on tissue quality, an assessment of RNA integrity was performed on bulk tissue preserved in RNAlater using independent RNA preparation methods at two different sites. All the bulk RNA samples (total 12, 6 nephrectomy samples in RNAlater shipped to each site) were sequenced at a central site. These results showed strong correlation among adjacent tissue samples from the same subject for all 6 subjects and established the shipping conditions for frozen tissue that do not adversely affect tissue state as measured by RNA expression and integrity analysis (**Fig. 5**). We noted several observations during the pilot experiment that could in general impact tissue integrity including insufficient dry-ice, the contents not well embedded in dry ice upon receipt due to movement during transit and frozen slides not secured in secondary box during transit (detailed shipping conditions are in the online pathology and biospecimen protocols at https://kpmp.org/researcher-resources/).
E. *Initial processing at the TISs.* This experiment also provided an opportunity to examine the initial processing steps at each TIS and explore the potential to standardize common procedures. This resulted in the implementation of common procedures at each site, which were incorporated in the KPMP TIS manual of procedures (www.KPMP.org). For example, this pilot experiment identified the need to obtain histology sections flanking areas of interrogation within the tissue, to inform on the state, composition, and orientation of the tissue. This process also allowed the same OCT block to be exchanged by two interrogation sites to perform successful molecular interrogation simultaneously with 3 different techniques (**Fig. 5**). The pilot experiment was also crucial to verify, validate and expand the metadata variables that needed to be captured for faithful documentation of the tissue journey from harvesting to interrogation.

**Fig. 5.**
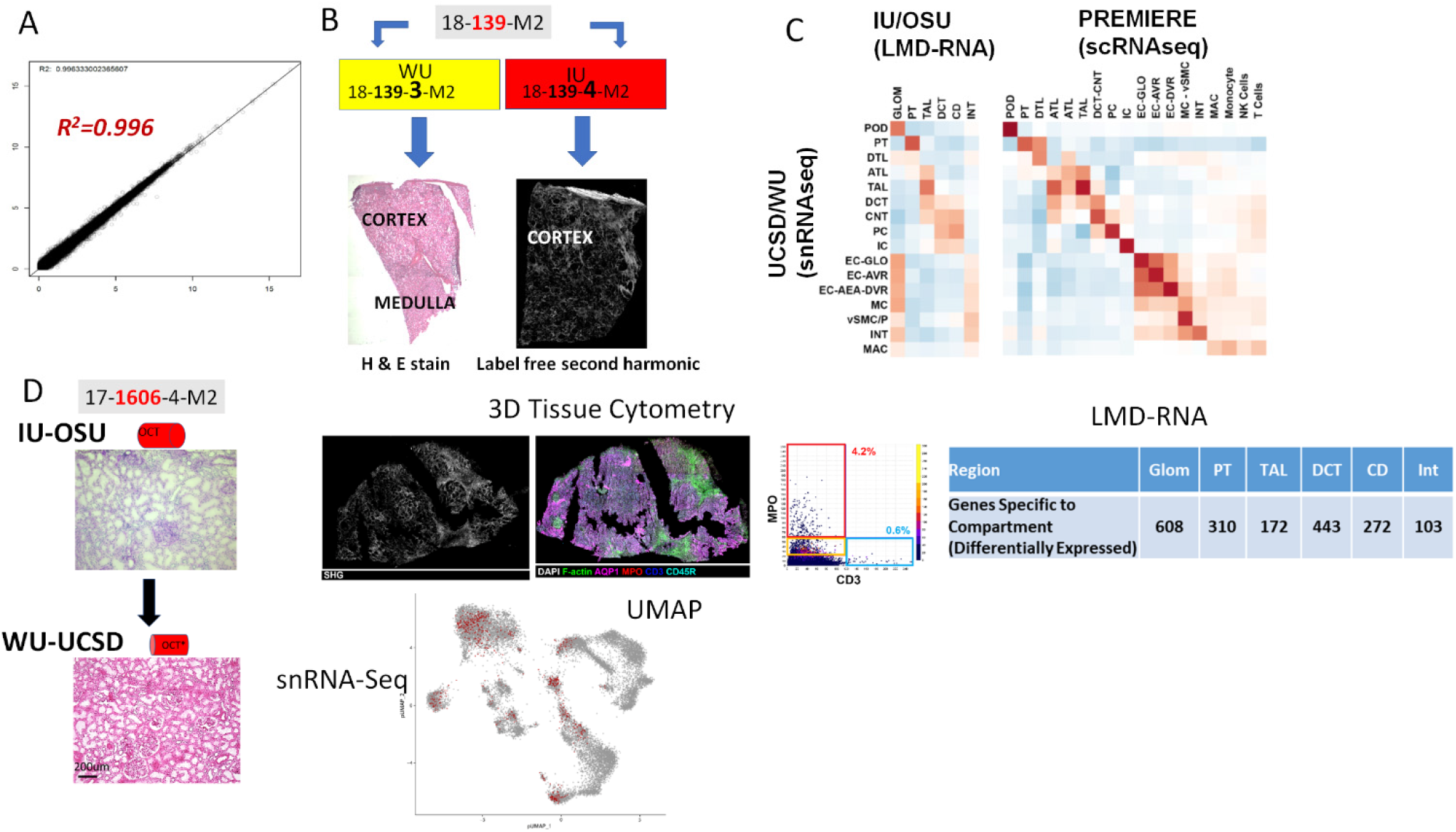
Testing the “Follow the tissue” pipeline with data generated from pilot samples and demonstrating feasibility of multiple technologies on limited tissue sample across sites. **A)** High correlation of bulk RNA seq from adjacent samples shipped and processed at different sites. **B)** Adjacent samples from same patient shipped to different TISs show different types of information from different technologies in KPMP with histological tissue composition verification for snRNAseq at WU-UCSD and label-free second harmonic generation highlighting extent of fibrosis at IU-OSU. **C)** Correlation between cell types and regions among snRNA-seq, LMD transcriptomics of the indicated regions by IU/OSU and between snRNA-seq and scRNA-seq. **D)** Economizing tissue usage by rotating fresh OCT frozen kidney tissue between IU and WU/UCSD. Here frozen sections were processed and analyzed for 3D tissue cytometry and LMD-RNA analysis and then the block was shipped to WU/UCSD for histological registration and subsequently generating snRNA-seq. Genes specific to each LMD compartment were determined by their upregulation in that compartment compared to other compartments as well as differential expression at a nominal p < 0.001. The UMAP plot shows the sample (~900 nuclei) contributing to many of the cortical cell types in the kidney snRNAseq atlas (~18000 nuclei).(11)

**Fig. 6.**
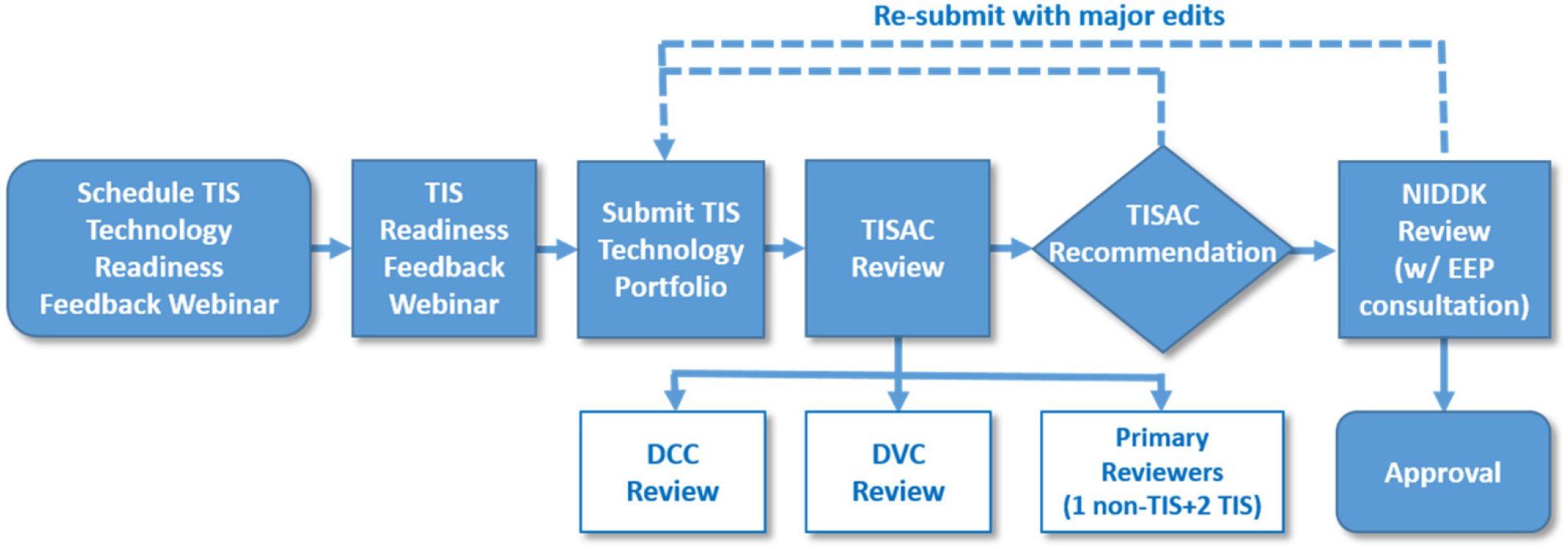
The tissue interrogation site approval committee (TISAC) workflow. This workflow includes completion of a recorded webinar and submission of a portfolio that details the quality control metrics and technology outcomes. The TISAC reviews this material, providing recommendations to the tissue interrogation site for improvement before a site is approved to receive biopsies. The TISAC ultimately forwards their recommendations to the NIDDK and external expert panel (EEP) for final approval. Data visualization center (DVC); data coordination center (DCC).

#### 2) Analytical QC parameters for each technology

One of the main goals of this experiment was to test and validate the QC parameters for each technology in different sites. The design of the pilot experiment allowed repeat testing on a single specimen to assess technical reproducibility and the use of tissue from different donors and tissue from different sources (pilot and local samples) ensured testing the methodologies for rigor and biological reproducibility. The QC parameters adopted by each technology based on this pilot experiment are summarized in **Supplemental Tables S1-3**.

Cross-validation with existing data/standards or by cross-validating outcomes from various technologies performed on the single source kidney tissue provided in this pilot provided another level of confirmation to the QC parameters. Examples of this include detection of the same molecules/metabolites in the same samples using different technologies and concordant “derived” readouts such as pathway analyses. For example, the TIS technologies can detect different genes/molecules/metabolites, but these molecular entities can be part of the same signaling pathway. Examples of orthogonal validation approaches are shown in **Fig. 5** and integrated analyses will be presented in a separate manuscript.

#### 3) Post-analytical outcomes

In addition to the cross-validation benefits discussed above, examining the outputs from various technologies promoted integration efforts and helped determine the extent of complementary information provided by each technology and further metadata harmonization at the various levels of tissue processing, analytics and analysis. Additionally, parameters for diagnostic features, composition, and integrity of the tissue that are applicable to all the TISs were further refined and led to a protocol for interrogating patient biopsies in the KPMP described in a comprehensive Pathology protocol document (https://kpmp.org/researcher-resources/)). This ensures that a comprehensive cellular and molecular converge is provided by the consortium to make a robust kidney atlas and a platform for discovery.

An additional important outcome was that significant amount of gene and protein expression data were generated from the pilot samples. These data collected from multiple sites provide an initial view of cellular diversity in the human kidney (**Table 3, Supplemental Fig. S3)**. The analysis also revealed stress states related to processing of tissue and underlying pathology that could not have been predicted from gross evaluations in presumably healthy tissue. In fact, some novel discoveries have already emerged in the initial version of the kidney atlas from the Pilot project.(11, 16)

**Table 3:**
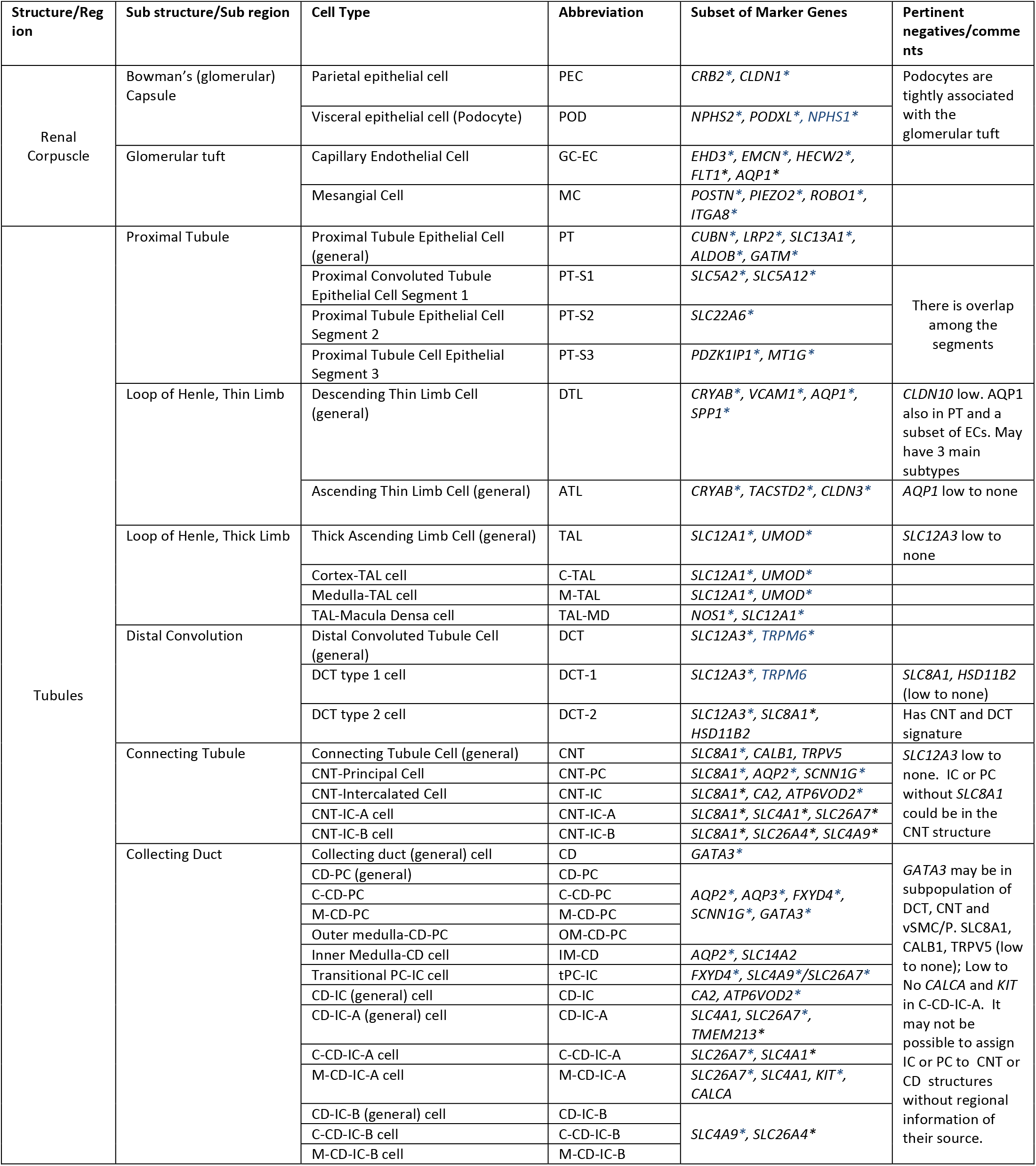
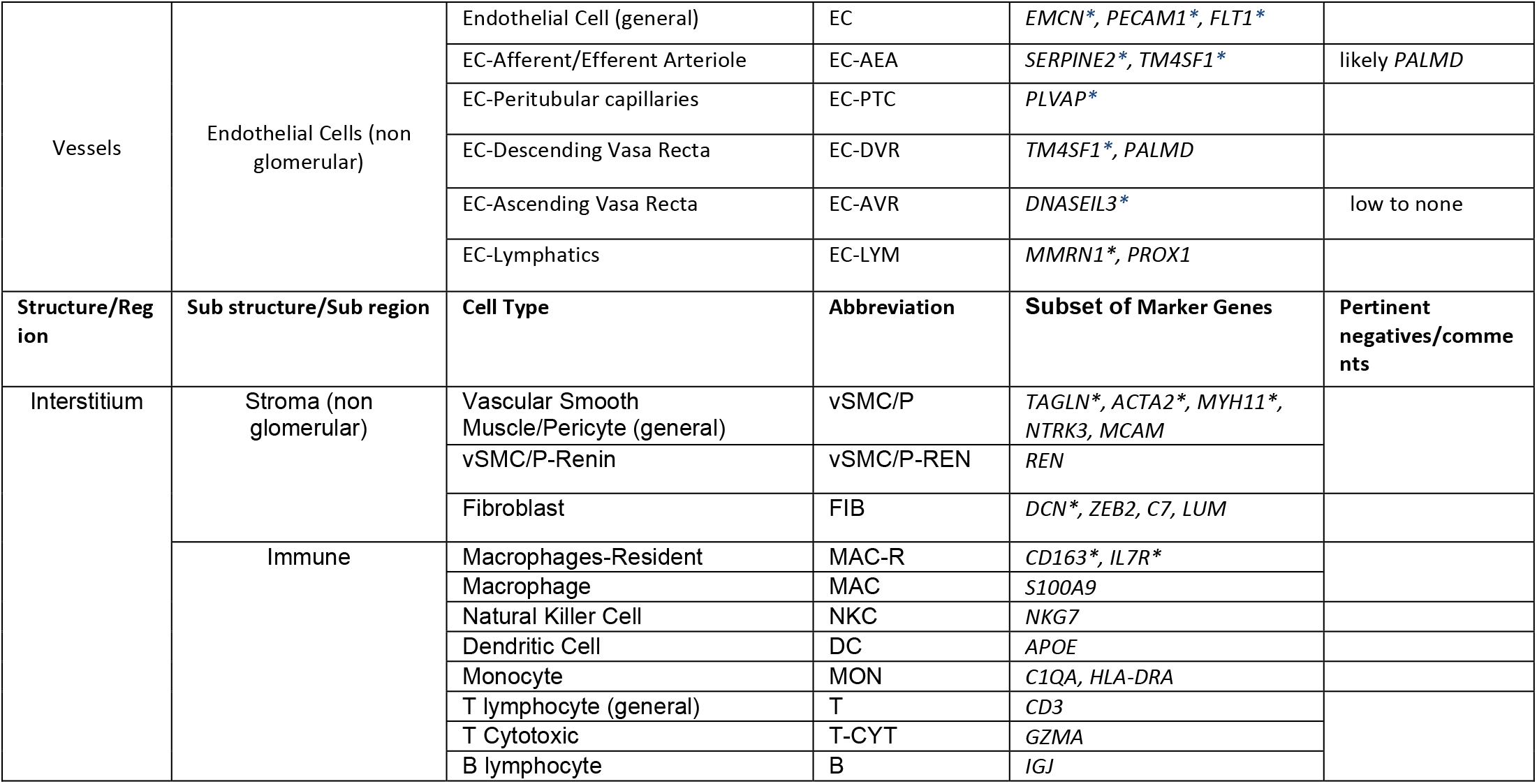
Cell types and associated markers from KPMP Pilot 1 transcriptomic studies. Asterisk denotes genes detected by more than one technology.

### Identification of gaps and improvement of the process

An area of priority identified during the integration efforts was the need to establish benchmarks for the nomenclature of cell types, regions and associated genes, proteins and metabolites for reference and disease atlas and various injury states. A promising methodology of analysis that could link multiple technologies is a cell-centric approach, whereby the outputs can reflect changes at the cell level in a tissue specimen. This was essential as several groups are investigating the single cell transcriptome or proteome of the kidney but there is lack of conformity regarding nomenclature and annotations. However, this analytical process requires an initial definition of cell types based on a set of criteria, such as: gene expression (RNA and protein), cell state (baseline, stress, injury), spatial localization and associations, among others. The pilot studies generated an initial working list delineating the complexity of cell types and a subset of associated marker genes in the adult human kidney which could serve as a starting point for kidney OMICS and imaging studies for classification of cell types and states and harmonization with recent renal tubule epithelial cell nomenclature (**Table 3**).(3) An ontological representation of the cell markers has been initiated to seamlessly link gene, cell type and spatial tissue location at an integrative semantic level.(7)

### Implementation of best practices to perform tissue interrogation on biopsies from the KPMP: the TISAC process

*Approval of TISs to receive biopsies for interrogation.* To rigorously evaluate each technology and eliminate self-approved bias by each TIS we established a Tissue Interrogation Site Approval Committee (TISAC). The committee is composed of representation across the KPMP, NIDDK and external *ad hoc* members as required to provide sufficient expertise to review the technology This committee evaluates all technologies prior to approval to perform studies on precious biopsy specimens and ensures the technology under consideration has presented evidence for robust QC metrics, sample handling, addressing batch effects, assay drift and is complementary with other technologies. These elements are summarized in **Fig. 3** and **Supplemental File 3**. The TISAC provides constructive feedback to the TIS whose technology is under review to enhance rigor, reproducibility and complementary aspects of each technology, as well as identify areas that require additional supporting data. Once satisfied with a given technology’s readiness, the TISAC recommends approval and notifies the Steering Committee. The TISAC provides their report to the NIDDK and the KPMP external expert panel who ultimately approve the technology and TIS for receipt of patient samples. Each TIS is further expected to report on the state of their technology at least annually and sooner if there are any modifications in the protocol used.

### Ongoing progress, challenges and future outlook

Evaluating progress and perceiving challenges require a continuous process of self-evaluation at several levels in the KPMP including the expertise of external investigators and interaction with large national and international consortia. The important components for a healthy and sustainable workflow for “follow the tissue” are: a) identifying opportunities to improve the quality control process, b) tackling challenges introduced by evolving or newer technologies, and c) mitigating potential threats or unforeseen errors. Examples of ongoing areas of development or challenges in the immediate future are described in **Table 4**.

**Table 4:** Challenges and opportunities ahead.

| Challenges | Explanations |
| --- | --- |
| Evolving ontology representation of various data and metadata elements | Generation of data and metadata will continue to grow. The need to link different and novel data types, cell markers, assays and assay components is a dynamic approach and need constant community engagement. |
| Data visualization, integration and dissemination of results | Data (raw and processed), metadata and all QC elements will become publicly available for all type of users in a way easy to find, access, interoperate, reusable and interpret. What is the best way to do this? |
| Incorporation of external data | The external data would need to meet KPMP QC standards for meaningful interpretation and relevance and reach of KPMP and nonKPMP generated data for discoveries |
| KPMP policies to address changes in technologies | Some technologies and platforms may change. How will data be acquired and archived to be compatible with data from future platforms? This may need a large source of reference tissue that can be interrogated and shared by the TISs. |
| Software changes (analytic or visual software) | New software present challenges in compatibility, reliability and security. |
| Strategies to test batch effect and technology or assay drifts | Current and future technologies are expected to provide an *a priori* plan to detect batch effect and technology drifts and provide solutions. What reference tissue standards are suitable for this purpose? |
| Validation of emerging technologies and incorporation into KPMP. | Will there be a need for a standard tissue used for validating new technologies? What should this tissue be? Is there unlimited supply? |
| Hyperdimensional data management, storage and sharing | An increasingly problematic issue when big data will be generated from each tissue specimen. |
| Patient protection in the era of artificial intelligence. | The risk of linking patients to de-identified raw data may increase as machine learning tools develop further. Steps to mitigate risks in publicly available data will need to be implemented. |
| Justifying the use of limited renal biopsy tissue for research that is unlikely to benefit the patient and could compromise diagnostic yield | The follow the tissue pipeline enables multimodal analysis on left over tissue from diagnostic specimens and would enhance current diagnostic pipeline as these technologies may lead to new validated clinical tests that can improve diagnosis and management of patients with kidney disease. |

One of the limitations of implementing novel high-throughput technologies on human biopsy samples is the lack of knowledge of assay variances and the biological variability between samples resulting from several factors (discussed above). This is further complicated by the complexity of cell types and 3D relationships which have never been explored with the resolution of the current scale of technologies. These factors pose a challenge in performing power calculations and estimating sample size for reference or kidney disease atlas. For example, the variance in mean gene expression could be different in distinct cell types depending on demographics or underlying pathology. It is likely that analyzing reference kidney and disease biopsy tissue from first 20 participants will provide insights into these variations and inform on the sample sizes needed for different cell types under different healthy or disease contexts. In the KPMP we will analyze results from 20, 50, 100 and 200 biopsies to get a better understanding for sample size needed for each of the disease categories. In addition, there is already an inherent sampling bias to derive conclusions because a biopsy represents only a fraction of the entire kidney. This limitation already exists in current clinical and pathological evaluations for any organ, however, the scale of multimodal analysis presented above provides a more rigorous analysis of the tissue and when enhanced by increased sample size, will likely overcome this limitation (Table 4).

## Conclusions

With the implementation of a standardized multimodal and integrated pipeline for molecular interrogation of kidney biopsy specimens, the goal of the KPMP is to set high standards for quality control, rigor and reproducibility. Vetted technologies participating in the KPMP will undergo careful scrutiny to comply with these goals of quality control, while at the same time allowing a dynamic and iterative approach that promote improvement and transparency. In doing so, the KPMP could become a model for other national and international efforts that also seek to decipher human disease and build a dynamic tissue atlas. With the QC infrastructure in place, the KPMP will achieve its goal to improve patient care and provide data to develop therapeutics for kidney disease with rigor and reproducibility.

## Author Contributions

MTE, RM, BBL, TS, TA, AS, SP, CRA, DD, EAO, SW, GZ, MJ and KD performed the ground work for the quality control group, wrote the KPMP TIS manual of procedures and generated figures. HH, JZ, RS, RM, TS, EAO, MTE, TME, PH, SP, MK, ZL and SJ generated the initial working reference marker list. CEA, TME, JBH, JL, MK and SJ led the Pilot 1 protocol. TME, VD, LB, JG, CEA, ZL, SJ and JBH developed the pathology QC tissue qualification and tissue processing criteria. JBH organized and executed the Pilot tissue collection and distribution. JC designed the SpecTrack system. CP prepared and organized the TIS manual of procedures and performed data organization services. YH led ontology development for QC metadata and knowledge standardization. BS and EA organized data integration efforts and data authentication in the data hub. KS and MS led the OMICS discussion group. SJ led the quality control group. TME and CEA led the tissue processing group. MTE and SJ led the Molecular and Pathology Integration group. MTE and SJ conceived and led the TISAC process. RI, OGT KZ, ZL, PH, BR, PCD, KS, MS, JBH, CEA, LB, JG, TME, MK and SJ conceived the integrated TIS pipeline and QC vision. TME and SJ wrote the initial draft of the paper. All authors contributed to the writing and editing of the manuscript.

## Acknowledgements

The Kidney Precision Medicine Project is supported by the National Institute of Diabetes and Digestive and Kidney Diseases through the following grants: UH3 DK114923, UH3 DK114920, UH3 DK114933, UH3 DK114937, UH3 DK114907, U2C DK114886. We thank the KPMP patient participants, scientific officers from the NIDDK, Recruitment sites, Central Hub and all the TISs for many valuable discussions and feedback towards the QC efforts. We are grateful to the KPMP Publications and Presentation committee for suggestions and review of this manuscript. A complete list of all KPMP members can be found at kpmp.org.

## Supplemental material table of contents

**Supplemental table S1.** Standardization and Quality Control Parameters for Transcriptomics Technologies.

**Supplemental table S2.** Standardization and Quality Control Parameters for Proteomics and Metabolomics Technologies.

**Supplemental table S3:** Label-based imaging QC.

**Supplementary table S4.** Harmonized imaging parameters and metadata to be recorded across TIS sites.

**Supplemental Figure S1:** Sectioning design for a same source tissue KPMP pilot

**Supplemental Figure S2:** Spectrack software and shipment workflow.

**Supplemental Figure S3:** Kidney regions and cell types.

**Supplemental File 1.** Overview of KPMP technologies-supplementary methods

**Supplemental File 2.** Initial candidate kidney marker panel

**Supplemental File 3.** Tissue interrogation site approval committee (TISAC) checklist.

