## Supplemental material for "A Multimodal and Integrated Approach to Interrogate Human Kidney Biopsies with Rigor and Reproducibility: The Kidney Precision Medicine Project"

**Supplemental material table of contents**

**Supplemental table 1.** Standardization and Quality Control Parameters for Transcriptomics Technologies.
**Supplemental table 2.** Standardization and Quality Control Parameters for Proteomics and Metabolomics Technologies.
**Supplemental table 3:** Label-based imaging QC.
**Supplementary table 4.** Harmonized imaging parameters and metadata to be recorded across TIS sites.

**Supplemental Figure 1:** Sectioning design for a same source tissue KPMP pilot
**Supplemental Figure 2:** Spectrack software and shipment workflow.
**Supplemental Figure 3:** Kidney regions and cell types.

**Supplemental File 1.** Overview of KPMP technologies- supplementary methods
**Supplemental File 2.** Initial candidate kidney marker panel
**Supplemental File 3.** Tissue interrogation site approval committee (TISAC) checklist.

**Supplemental table 1. Standardization and Quality Control Parameters for Transcriptomics Technologies.** "*" denotes technologies approved by TISAC for interrogating patient biopsies (Jan 2020).

| **Category/Site** | **UCSD-WU** | **PREMIERE** | **UCSF** | **IU-OSU** | **Standardized QC** |
|---|---|---|---|---|---|
| **Technology** | **snRNA-seq*** | **scRNA-seq*** | **mDroscRNA-seq** | **LMD mRNA* subsegmental and bulk miRNA** | **All RNA OMIC** |
| **Sample Preservation/Storage/Processing/ Transport Parameters** | • Fresh frozen OCT<br>• -80 °C<br>• Cryosection (2-20mm$^3$), RNAlater<br>• Nuclear prep (>10K)<br>• Dry ice | • Fresh frozen Cryostor<br>• -80 °C<br>• Dissociation (Liberase)<br>• Dry ice | • Fresh frozen Cryostor<br>• -80 °C<br>• Dissociation (Liberase)<br>• Dry ice | • Fresh frozen OCT<br>• -80 °C<br>• mRNA: Cryosection, dissect subsegments < 2 h, >500Ksqm<br>• miRNA: Cryosection 20-60 µm<br>• Dry ice | • Pre-analytical<br>• Tissue procurement<br>• Tissue processing (preservation, path assessment)<br>• Shipping (dry ice)<br>• Supplies<br>• Storage (-80 °C) |
| **Assay (RNA QC, library preps, Instrument, Sequencing)** | • Lib size (200-1000bp)<br>• Bioanalyzer 2100<br>• 10X Chromium<br>• Illumina<br>• Paired end (30x100) | • Lib size (200-1000bp)<br>• Bioanalyzer 2100<br>• Illumina<br>• 10X Chromium<br>• Paired end (30x100) | • Lib size (200-1000bp)<br>• Bioanalyzer 2100<br>• 10X Chromium<br>• Illumina<br>• Paired end (30x100) | • DV200 > 25%<br>• Lib size (200-1000bp)<br>• Bioanalyzer 2100<br>• RNA ref std (Stratagene)<br>• Illumina<br>• Paired end (30x100) | • RNA quality (DV200>25%; RIN for bulk > 6.0)<br>• Library size (200-1000bp)<br>• RNA ref std for bulk (Stratagene)<br>• Instrument<br>• Sequencing platforms |
| **Analytics (QC filters, artifacts, softwares, thresholds)** | • >400<7500 genes/nucleus<br>• Seurat V3 or Pagoda 2<br>• UMAP cluster<br>• Reference markers for annotations<br>• >30 cell/integrated clusters<br>• >100 QC nuclei/sample<br>• GRCH38 | • >500<5000 genes/cell<br>• MT<20% or <50% (Seurat V2/V3)<br>• UMAP cluster<br>• Reference markers for annotations<br>• >30 cell/integrated clusters<br>• >100 QC cell/sample<br>• GRCH38 | • >500<5000 genes/cell<br>• MT <50% (Seurat V3)<br>• UMAP cluster<br>• Reference markers for annotations<br>• >30 cell/integrated clusters<br>• GRCH38<br>• >100 QC cell/sample<br>• GRCH38 | • Read count > 10 over 50% of samples for at least 1 subsegment<br>• >1 million reads/sample<br>• Edge R<br>• Reference markers for annotations<br>• GRCH38 | • >400/500<5000 genes/cell (sn/sc)<br>• Seurat V3<br>• Reference genome GRCH38<br>• UMAP cluster<br>• Reference markers for annotations<br>• >30 cell/integrated clusters<br>• >50/100 (sn/sc) datasets per sample |
| **Data deposition** | • Fastq, bam<br>• Count matrix R Objects | • Fastq, bam<br>• Count matrix R Objects | • Fastq, bam<br>• Count matrix R Objects | • Fastq, bam<br>• Count matrix R Objects, csv, txt | • Fastq, bam<br>• Count matrix R Objects<br>• csv, txt |

**Supplemental table 2. Standardization and Quality Control Parameters for Proteomics and Metabolomics Technologies.** "*" denotes technologies approved by TISAC for interrogating patient biopsies (Jan 2020).

| **Category/Site** | **UCSF** | **IU-OSU** | **UTHSA-PNNL-EMBL** | **Standardized QC** |
|---|---|---|---|---|
| **Technologies** | **nscProteomics** | **LMD regional proteomics*** | **Spatial Metabolomics** | **All Proteomics and Metabolomics** |
| **Sample Preservation/Storage /Processing/ Transport Parameters** | ● Fresh frozen OCT<br>● -80 °C<br>● Cryosections<br>● Dry ice | ● Fresh frozen OCT<br>● -80 °C<br>● Cryosections<br>● Dry ice | ● LN2 snap frozen<br>● -80 °C<br>● Cryosection<br>● Dry ice | ● Pre-analytical<br>● Tissue procurement<br>● Tissue processing (preservation, path assessment)<br>● Shipping (dry ice)<br>● Supplies<br>● Storage (-80 °C) |
| **Assay** | ● adjacent sections > 10 cells per region<br>● Orbitrap Fusion (< 3 ppm calibration accuracy)<br>● Peptide standards | ● 10,000-20,000 glomerular or tubular cells<br>● Orbitrap Fusion (< 3 ppm calibration accuracy)<br>● Peptide standards | ● Entire tissue section, bulk analysis on remaining<br>● Orbitrap QE and FTICR-MS (< 3 ppm calibration accuracy)<br>● Commercial tune mix | ● Tissue sectioning<br>● Cell /region isolation<br>● Orbitrap Fusion (proteomics)<br>● Peptide standards (proteomics) |
| **Analytics** | ● >2000 proteins per 10 cell prep<br>● > 6aa<br>● <4600Da peptide mass<br>● Missed cleavage for peptides = 2<br>● < 1% FDR<br>● > 97% correlation for peptides in replicates<br>● Reference markers for annotations | ● > 3000 proteins identified<br>● < 1% FDR<br>● Reference markers for annotations | ● >100 annotated metabolites per section (<20% FDR)<br>● Detection of standard metabolites<br>● Detection of location specific metabolites<br>● Histology correlation | ● > 2000/3000 proteins (nsc/LMD)<br>● < 1% FDR (nsc, LMD)<br>● Standard metabolites and renal metabolites |
| **Data deposition (file types)** | ● .raw, .csv | ● .raw, .csv | ● MALDI-MSI: .d/.mis, .xml, .imzML, .ibd, raw, csv<br>● LC-MS/MS: .raw, .mzML | ● raw, csv (proteomics)<br>● raw (LC-MS/MS)<br>● d/mis, imzML, ibd, raw, csv (MALDI-MSI) |

**Supplemental table 3: Label-based imaging QC.** "*" denotes technologies approved by TISAC for interrogating patient biopsies (Jan. 2020).

| | **IU-OSU** | **UCSD-WU** | **UM-Broad-Princeton** | **UCSF** | |
|---|---|---|---|---|---|
| **Technology** | 3D Tissue Cytometry* | DART-FISH2 | ISH | mIFISH | CODEX |
| **Probe Targets Per Section** | 8 Immunofluor targets | >50 mRNAs | 3-10 mRNA | 20 mRNA & proteins | 30 proteins |
| **2D or 3D Imaging?** | 3D | 3D | 2D | 2D | |
| **Spatial Resolution** | 0.5-1 µm | 1 µm | single cell | 1µm/single cell | |
| **Cell type and structure mapping** | Validated antibodies and fluorescent small molecules known to bind specific cell types | nuclear stain, lectin labeling for PT or CD, autofluorescence | cell specific RNA and histological stain in sequential plane | Anchor (structural) and immune cell and interstitial compartment mapping at single cell resolution with direct H&E correlation | |
| **Area/Volume of Imaging Region** | Entire biopsy length (1-1.5cm) x 50µm thickness | 10-20 micron sections of 1 $mm^2$ | Entire biopsy length (1-1.5cm) x 5 µm thickness | Entire biopsy (20-25 $mm^2$) | |
| **Dimensions of Image Voxels** | 0.5 x 1µm x 1µm | 0.15 x 0.15 x 0.3 µm | N.A. | N.A. | |
| **Amount of Tissue** | Minimum: One 50 um thick section (imaging/staining) | 10 x 10 µm thick sections | 5 µm thick section for each probe target | One 3µm-thick FFPE section for each 21-plex mIFISH stain. Two 3µm-thick FFPE sections for a positive/negative ISH control. | One 5µm-thick OCT section for each 32-plex CODEX stain |
| **Pre-imaging QC** | -Primary Ab validation<br>-Pre-staining with secondary antibodies as a negative control<br>-AF/SHG (tissue quality)<br>-16 channel pre-stain imaging | -Validation of generated probes<br>-Maintenance of fluorescent imaging system | In development | -Instrument: Auto-calibrated with each use<br>-Tissue: H&E to assess tissue integrity<br>-Antibody validation: IPOX/IIF/orthogonal | -Instrument: standardized imaging of IF calibration slides at regular intervals<br>-Tissue: H&E-stained sections to assess tissue integrity/freezing artifacts<br>-Oligo-labeling: comparison of the staining across CODEX, indirect IF, and IPOX platforms |
| **Acquisition QC** | -Reuse acquisition settings<br>-Point Spread Function<br>-Signal to noise ratio | -Signal to Noise Ratio<br>-Number of rolonies decoded<br>-Number of genes decoded | -Negative control: DapB<br>-Positive control: housekeeping genes | -On-slide tissue microarray controls<br>-Built in software function to set exposure times<br>-New antibody lots compared to "gold standard" -- epifluorescent images for batch consistency<br>-ISH: positive & negative control probes | -Labeling-detection: on-slide tissue microarray controls<br>-Imaging: saturation based on histogram<br>-Cell clustering by intensity gating: visual assessment of the raw image |
| **Analysis QC** | Unmixing (labeled beads at known ratios)<br>Segmentation quality (F1 scores) | -Misclassification rate (accuracy of barcode decoding)<br>-Number of rolonies per cell | In development | Visual inspection of analyzed images: segmentation & analyte detection accuracy | Cell clustering by intensity gating & visual assessment of the raw image |
| **Imaging and Analysis Throughput** | 2-3 samples per week | 1 section per week | In development | 8 sections/week (mIFISH) | 2-3 sections/week (CODEX) |

**Supplementary table 4. Harmonized imaging parameters and metadata to be recorded across TIS sites.**

| | | Feature | Value (ex. IU/OSU) |
|---|---|---|---|
| **Sample characteristics** | | Specimen ID | **18-142-04** |
| | | Derived specimen ID | **18-0007** |
| | | Data Lake Package ID | 9d401b61-55f2-4d8b-93f9-f473756d9cfc |
| | | Tissue thickness | 50 microns |
| | | Gross sample quality | Annotated macro image |
| | | Quantified sample quality | RNA quality (DV200 value) |
| | | Sample preparation - methods | MOP version number |
| | | Sample preparation - reagents | Vendor, catalog number, batch |
| | | Image sample preparation - quality control | Reagent validation images |
| **Image data and collection** | **Raw target image data** | Technology | 3D multiple fluorescence |
| | | Number of targets | 8 |
| | | Target IDs | THP, F-actin, AQP1, CD3,CD68, SIGLEC8, MPO, DNA |
| | | Sample volume imaged | Entire area (e.g., 3 mm x 6 mm) x 50 microns depth |
| | | Voxel dimensions | 0.5 x 0.5 x 1 micron |
| | | Raw target image file format | 16 channel Leica .lif |
| | | Raw target image file size | 100 GB |
| | | Image acquisition metadata | Leica lif file |
| | **Raw label-free image data** | Technology | Autofluorescence, SHG |
| | | Sample volume imaged | Entire area (e.g., 3 mm x 6 mm) x 50 microns depth |
| | | Voxel dimensions | 0.5 x 0.5 x 1 micron |
| | | Raw label-free image format | 2 channel Leica .lif |
| | | Raw label-free image file size | 10 GB |
| | | Image acquisition metadata | Leica lif file |
| | **Image collection** | Image collection - methods | MOP version number |
| | | Image collection - quality control | Image of standard beads (SNR) |
| | | | Image of resolution standard beads (PSF) |
| **Image analysis** | | Image analysis methods | MOP version number |
| | | Image analysis software | VTEA 0.5.2 |
| | | Image processing record | VTEA log file |
| | | Image processing QC metrics | Spectral deconvolution of beads |
| **Results** | | Output, deconvolved target image data | 3D, 8-channel fluorescence image volume |
| | | Output target image file format | .tiff |
| | | Output target image file size | 50 GB |
| | | Output label-free image data | 3D, 2-channel autofluorescence/SHG image |
| | | Output label-free image format | .tiff |
| | | Output label-free image file size | 10 GB |
| | | Image visualization access | Online 3D rendering |
| | | Derived measurements | Total cell number. Numbers of PT, TAL, vascular endothelia, T-cell, macrophages, neutrophils, eosinophils |
| | | Measurement data files | Raw results |
| | | | Supervised data analysis |
| | | | Unsupervised data analysis |
| | | Data visualization access | VTEA |
| | | Image cross-mapping | PAS in adjacent section |
| | | Omics cross-mapping | RNA from LMD in next section |

### Supplementary figure 1

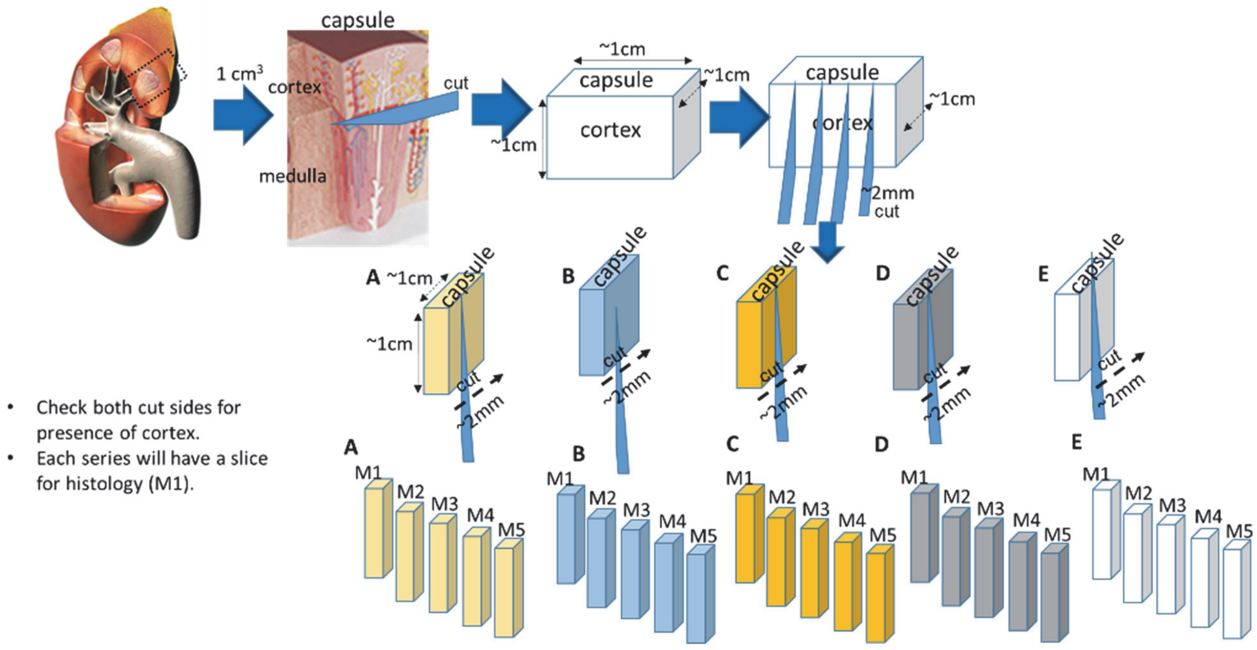


**Supplemental Figure 1: Sectioning design for a same source tissue KPMP pilot**

### Supplementary figure 2

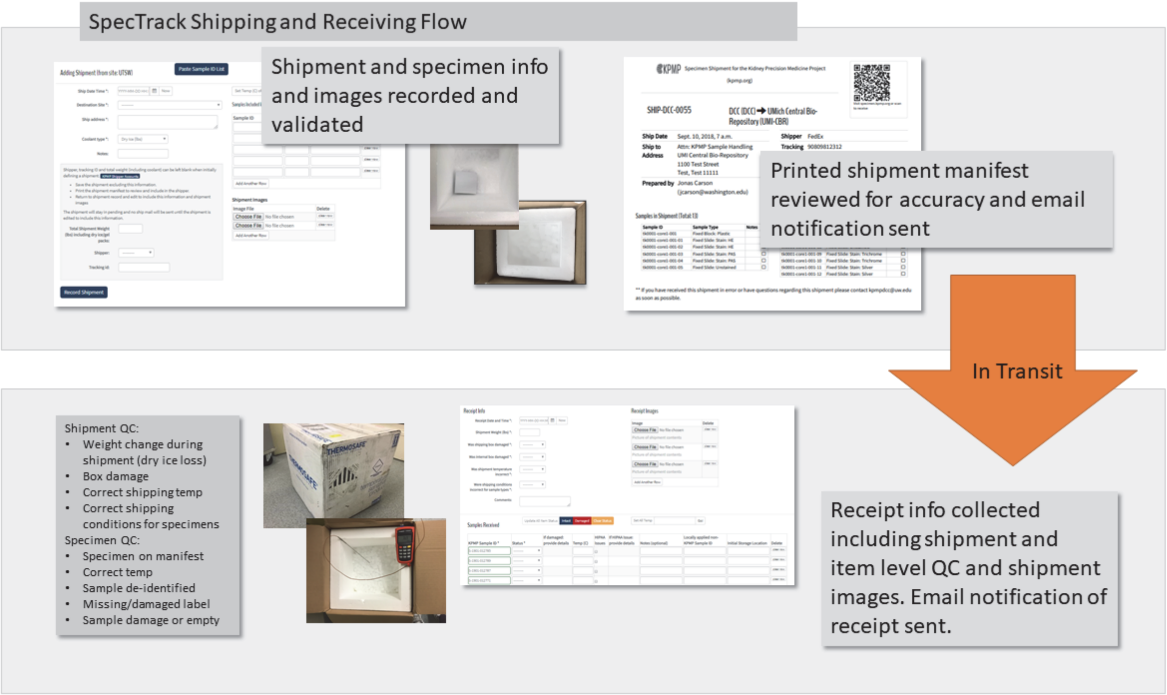


**Supplemental Figure 2: SpecTrack software and shipment workflow**. Spectrak is a customized software designed by the KPMP to track in real-time the tissue transfer and all quality control metrics associated with shipment and handling.

### Supplementary figure 3

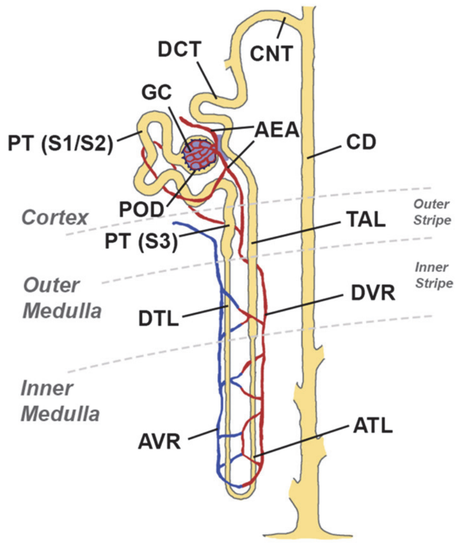


**Renal Corpuscle (Bowmans capsule and Glomerulus)**
*Parietal epithelial cells, Podocytes (POD), Mesangial cells (MC), Endothelial (EC)*

**Renal Tubule**

***Proximal Tubule Epithelial Cells***
*Proximal Convoluted (PT S1/2)*
*Proximal Straight (PT S3)*

***Distal Convoluted Tubule Cells***
*Distal Convoluted Tubule (DCT1)*
*Distal Convoluted Tubule (DCT2)*

***Loop of Henle Cells***
*Descending Thin Limb (DTL)*
*Ascending Thin Limb (ATL1-3)*
*Thick Ascending Limb (TAL1-3)*

***Collecting System Cells***
*Connecting tubule (CNT, PC, IC-A, IC-B) (Cortex)*
*Collecting Duct (PC, IC-A, IC-B, IMCD) (Cortex, Outer/Inner Medulla)*

**Interstitium Cells (extra glomerular)**
Endothelial (1-4), vSMC/Pericytes, Fibroblasts, Immune Cells (T, B, NK, MAC, Dendritic)

**Supplemental Figure 3: Kidney regions and cell types.** The nephron illustration is based on Kriz, W. & Bankir, L. A standard nomenclature for structures of the kidney. The Renal Commission of the International Union of Physiological Sciences(IUPS).Kidney Int.33,1–7 (1988). T, T lymphocyte; B, B lymphocyte; NK, natural killer cell; MAC, macrophage.

**Supplemental File 1.** Overview of KPMP technologies- supplementary methods

*Transcriptomics.* Several technologies are generating gene expression data at bulk, regional and single cell level for comprehensive coverage of the transcriptome for the reference (relatively healthy or pathologically normal) and disease atlases (AKI and CKD subtypes). Single nucleus (sn) (UCSD/WU) and single cell (sc) (UCSF and Premiere -Michigan/Broad/Princeton sites) RNA-seq technologies are being used for cataloging molecularly-defined cell populations for the generation a single cell kidney atlas.[1-3] Each of these technologies uses different approaches in sample preparation, dissociation or processing that have unique advantages and disadvantages (see TIS protocol on https://kpmp.org/researcher-resources/). Given the potential to introduce artifacts or miss cell types due to processing of these specimens, concurrent bulk RNA expression analysis on undissociated tissue or regional laser microdissection (LMD) (IU/OSU) is performed.[4-6] Using the LMD technology, transcriptomic signatures are identified for seven sub-segments of the nephron including the proximal tubule (PT), thick ascending loop of Henle (TAL), distal convoluted tubule (DCT), collecting duct (CD), as well as compartment-specific signatures for glomeruli, the tubulo-interstitium (TI), the interstitium (without glomeruli or tubules), as well as a bulk cross-section of the entire biopsy. LMD transcriptomic signatures can serve as an important independent validation measure that provides regional spatial context to cell populations discovered using single cell technologies. To complement the mRNA signatures obtained from the single event technologies and regional transcriptomics, miRNA sequencing of bulk cross-sections from the same OCT (optimum cutting temperature embedding media)-embedded core are sequenced specifically for small RNA (IU/OSU).[7]

*Proteomics.* Two different approaches were planned in the KPMP to generate reference and diseased kidney proteome. The regional/segmental approach by the IU/OSU group combines the collection of specific regions of the kidney using LMD with quantitative proteome analysis using state-of-the-art high performance liquid chromatography (HPLC)/mass spectrometry (MS) instrumentation to generate agnostic global profiles of the glomerular and tubulo-interstitial compartments with high sensitivity and reproducibility.[8] The UCSF group is employing recently developed nanoscale proteomics analysis in which a few cells can be processed to generate cell-specific protein profiles.[9] Combining this technology with regions isolated by LMD provides regional and spatial definitions of generated proteome data. Both technologies additionally provide bulk proteomics data on tissue sections allowing cross platform/site analysis.

*Metabolomics.* The UTHSA/PNNL group will generate spatial metabolomics measurements by using matrix-assisted laser desorption/ionization (MALDI) mass spectrometry imaging (MSI), an approach that has been utilized previously for kidney molecular imaging.[10, 11] Here, fresh-frozen tissues are cryosectioned, mounted on optically transparent and electrically conductive slides, coated with an organic matrix that assists in facilitating desorption and ionization of endogenous molecules, and serially probed with a laser to attain mass spectral information at predefined locations. They employ two different platforms to generate metabolite profiles that are designed for cross validation and complementarity. An important aspect is that data are processed using the METASPACE platform (a tool developed by the EMBL portion of this group),[12, 13] which is an automated molecular annotation engine that enables data visualization and co-registration with other optical images.

*2D and 3D Imaging*. There are several imaging technologies within KPMP that inform on precise 2D and 3D expression relationships of biomolecules, cells and structures with varying degree of multiplexing, spatial resolution and tissue preservation conditions **(Fig. 2)**. The UCSD/WU group will develop Decoding Amplified taRgeted Transcripts with Fluorescence In-Situ Hybridization (DART-FISH) to delineate single cell mRNA expression of several hundred transcripts in each cell at high resolution in 3D using high resolution confocal microscopy. The UCSF and PREMIERE sites will use miFISH to generate simultaneous gene and protein expression data at the single cell level in 2D space.[14] The IU/OSU site will utilize large scale 3D confocal imaging of longitudinal kidney thick sections probing concurrently for 8 different targets using antibodies or fluorescent small molecules, followed by tissue cytometry analysis.[15, 16] Prior to any staining, this technology uses label-free imaging to determine tissue integrity using endogenous fluorescence and quantify collagen deposition using second harmonic generation. The resulting image volumes undergo cytometry analysis with a customized Volumetric Tissue Cytometry and Analysis (VTEA) software. This approach allows the interactive exploration of the image volumes, as well as quantitative analysis of the abundance, distribution, and other 3D spatial features of interest for various cell types, based on supervised and unsupervised analytical approaches.[17] The UCSF site will use Co-detection by indexing (CODEX),[18] an orthogonal approach to perform 2D profiling *in situ* on up to 30 antigens at single-cell resolution on a single tissue section using antibodies labeled with unique oligonucleotide tags (“barcodes”). The labeling and detection is done in an iterative manner in groups of two or three targets per cycle by fluorescent dyes labeled with oligonucleotide sequences (“reporters”) corresponding to a given subset of antibody barcodes; in each cycle the previous reporters are removed and a new set is introduced. The montage image consisting of all signals from each cycle is analyzed for marker intensity distribution across the tissue section, for cell count of a given cell population defined by marker sets and for spatial relationship between cell populations of interest in the specimen.[18]

**Supplemental File 2.** Initial candidate kidney marker panel

See attached Excel spreadsheet

**Supplemental File 3.** Tissue interrogation site approval committee (TISAC) checklist.

**Kidney Precision Medicine Project**

### Tissue Interrogation Site (TIS) Technology Readiness Checklist

**Date:**

**Tissue Interrogation Site:** ______________________ **Technology:** ______________________

**Investigator Contact:** ______________________ E-mail: ______________________

Investigator: ______________________ E-mail: ______________________

Investigator: ______________________ E-mail: ______________________

| **A. TIS technology information (<u>COMPLETED BY TIS</u>)** | Pg. # |
|---|---|
| 1. Number of samples tested, central and local, that demonstrate feasibility. | _____ |
| 2. TIS MOP *Version*: _____ | _____ |
| 3. Sample sources (type and quantity; ex: 6 University of Michigan tumor nephrectomy pilot samples) (*include as many sample sources as necessary – bullet points below can be copied and pasted if more than 3).<br>• # Samples Source 1: _____<br>• # Samples Source 2: _____<br>• # Samples Source 3: _____ | _____ |
| 4. Technology data presented on TIS technology readiness feedback webinar (required), online calls, or at face-to-face meeting (specify below).<br>• _____<br>• _____<br>• _____ | _____ |

| **B. Central Hub / DVC Approval (<u>COMPLETED BY CENTRAL HUB/DVC REVIEWERS</u>)** | Pg. # | Yes | No |
|---|---|---|---|
| 1. Sample Handling (Central Hub approver: _____) | | | |
| • Demonstrate satisfactory use of specimen tracking software | _____ | ☐ | ☐ |
| • Verification of shipped samples (*include images in SpekTrack*) | _____ | ☐ | ☐ |
| 2. Detailed QC criteria for technology identified and included in TIS MOP. | _____ | ☐ | ☐ |
| 3. Detailed protocol included in TIS MOP and deposited in Data Lake. | _____ | ☐ | ☐ |
| 4. Demonstrated satisfactory collection and submission of metadata for sample processing as relevant to the technology (DVC approver: _____) | _____ | ☐ | ☐ |
| 5. Submitted data and associated metadata that is robust for integrative analysis (DVC approver: _____) | _____ | ☐ | ☐ |

**C. Central Hub / DVC Approval additional comments and recommendation section**

**Kidney Precision Medicine Project**

### Tissue Interrogation Site (TIS) Technology Readiness Checklist

| D. TIS technology readiness (COMPLETED BY TISAC REVIEWER: ______) | Pg. # | Yes | No |
|---|---|---|---|
| **1. Feasibility and Validation (Rigor and Reproducibility)** | | | |
| a. Submitted evidence that technology is validated on multiple samples from different sample sources. | _____ | ☐ | ☐ |
| b. Reported success-failure rates per the QC metrics that are included in the TIS MOP. | _____ | ☐ | ☐ |
| c. Provided supporting data to demonstrate feasibility, as well as technical and biological reproducibility (reviewer provide detailed input in section E). | _____ | ☐ | ☐ |
| d. Presented data to show local analytics of technology are robust (such as rigor used in identifying cell types or structures with associated genes/proteins/metabolites, clusters, artifacts in data, image analysis). | _____ | ☐ | ☐ |
| e. Provided supporting orthogonal validation data such as from knowledge base, other methods being used in TIS using local or pilot data that lend confidence to the results. | _____ | ☐ | ☐ |
| f. Provide supporting data to show how batch effects are addressed experimentally or analytically. | _____ | ☐ | ☐ |
| g. Proposed plans for assessing performance of technology if there is assay drift or changes arise in SOP, reagents, instruments, or analytics. | _____ | ☐ | ☐ |
| h. Provided supporting data that technology works on an amount of tissue similar to a biopsy. | _____ | ☐ | ☐ |
| **2. Technology Outcomes** | | | |
| a. Identified strengths of technology with supporting data. | _____ | ☐ | ☐ |
| b. Identified limitations of technology from the analysis done. | _____ | ☐ | ☐ |
| c. Provided justification for how technology is significant for KPMP, especially for building a reference and disease kidney atlas that will be used for patient management ultimately. | _____ | ☐ | ☐ |
| d. Provided data supporting a complementary role or unique contribution of this technology to the KPMP consortium. | _____ | ☐ | ☐ |

**E. TISAC reviewers' additional comments and recommendation section. Include overall assessment of technology:**

- **What are the conclusions?**
- **Do the strengths outweigh the limitations?**
- **Can the limitations be addressed with further research?**
- **Should this technology be prioritized?**
- **Does the site provide data supporting a complementary role or unique contribution of this technology to the KPMP consortium?**

### Tissue Interrogation Site (TIS)
### Technology Readiness Checklist

| F. Tissue Interrogation Site Approval Committee (TISAC) Approval Workflow (COMPLETED BY TISAC CHAIR: _____) | Yes | No |
|---|---|---|
| 1. Technology portfolio assigned and reviewed by three primary reviewers (two TIS and one non-TIS). Can solicit ad-hoc members from KPMP or externally if expertise is needed.<br>• TIS Reviewer 1: _____<br>• TIS Reviewer 2: _____<br>• Non-TIS Reviewer 3: _____ | ☐ | ☐ |
| 2. Portfolio containing supporting documentation (in .pdf format) of technology readiness criteria including the checklist received by TISAC (links to raw or analyzed data can be included). | ☐ | ☐ |
| 3. Technology portfolio including Reviewer comments electronically shared for review with entire TISAC. | ☐ | ☐ |
| 4. Technology portfolio discussed and reviewed on TISAC review web conference call.<br>• Date of web conference call: _____ | ☐ | ☐ |
| 5. TISAC formulates recommendation for technology portfolio:<br>• ☐ Accept<br>• ☐ Accept pending minor revisions<br>• ☐ Re-submit with major edits | ☐ | ☐ |
| 6. If accepted, technology portfolio reviewed and approved by NIDDK in consultation with External Expert Panel (EEP). | ☐ | ☐ |
| 7. TISAC provide composite report including NIH/EEP recommendations to the TIS and recommendations for re-assessment (e.g. modifications to technology, # of samples processed, etc.). | ☐ | ☐ |
| 8. Upon satisfactory demonstrating that all readiness criteria have been met and addressed all critiques with no major concerns, TISAC approves the technology and notifies Steering Committee. | ☐ | ☐ |

**G. TISAC additional comments and recommendation section**

**H. Summary and Recommendation, including timeframe for technology re-assessment**
